# The Origins of ATP Dependence in Biological Nitrogen Fixation

**DOI:** 10.1101/2024.02.22.581614

**Authors:** Derek F. Harris, Holly R. Rucker, Amanda K. Garcia, Zhi-Yong Yang, Scott D. Chang, Hannah Feinsilber, Betül Kaçar, Lance C. Seefeldt

## Abstract

Life depends on a conserved set of chemical energy currencies that are relics of early biochemistry. One of these is ATP, a molecule that, when paired with a divalent metal ion such as Mg^2+^, can be hydrolyzed to support numerous cellular and molecular processes. Despite its centrality to extant biochemistry, it is unclear whether ATP supported the function of ancient enzymes. We investigate the evolutionary necessity of ATP by experimentally reconstructing an ancestral variant of the key N_2_-reducing enzyme nitrogenase. We show that the ancestor has a strict requirement for ATP and its hydrolysis is coupled to electron transfer for N_2_ reduction. Our results provide direct laboratory evidence of ATP usage by an ancient enzyme, and underscore how biomolecular constraints can entirely decouple cofactor selection from environmental availability.

## Introduction

The use of ATP as an energy carrier is a ubiquitous feature of life. Energy transfer reactions that depend on the hydrolysis of ATP to ADP (typically complexed with Mg^2+^) and Pi are shared by all modern organisms. This universality and its role in RNA synthesis suggests that ATP is a remnant of ancient, possibly prebiotic biochemistry.^1,2^ Indeed, extant enzymes that require or synthesize ATP are similarly widespread across the tree of life and are proposed to have originated prior to the Last Universal Common Ancestor.^3–5^

The role of ATP in biology is not only ancient, but over more than 3.5 billion years of evolution,^6^ it emerged as the principal energy currency of life.^1,7^ It is not known when or why the predominance of ATP relative to other nucleoside triphosphates, like GTP, was established as the energy carrier. Several lines of evidence suggest that the landscape of energy carrying molecules in ancient cells may not have mirrored that of their modern descendants. Evolutionary switches in the ATP or GTP specificities of enzymes are known to have occurred, for example in the early-evolved family of P-loop NTPases.^8–10^ Further, experimental studies have demonstrated that such specificities can be quite mutable, mediated by even a single amino acid substitution.^7,11^ Finally, though modern ATP-(or GTP-) binding enzymes can require high specificity for their cognate nucleotide,^12,13^ many do exhibit promiscuous behavior and retain a lower level of activity with the alternate nucleotide^.10,14,15^ More broadly, substrate promiscuity has been observed in a number of reconstructed, phylogenetically inferred ancestral enzymes,^16^ though promiscuity as an inherent property of ancestral enzymes has been debated.^17,18^ Regardless of the general trend, these findings together raise the possibility that ancestors of ATP-dependent enzymes may not have exhibited the same specificities, and thus the requirement for ATP, as life’s primary energy currency may not have been as stringent for ancient cellular organisms.

Here, we investigate this possibility by examining enzymatic ATP requirements in the evolutionary history of a critical microbial metabolism: biological nitrogen fixation. Life on Earth requires a sustained availability of fixed N, an essential element of many biological molecules. However, throughout Earth’s history, it is thought that surface N has existed primarily in the form of atmospheric dinitrogen (N_2_),^19^ which is not directly usable for incorporation into biomolecules. To be biologically available, N_2_ must be fixed by reduction to ammonia (NH_3_) or oxidation to nitrogen oxides (NO_x_).^20–22^ There is evidence from the geologic record for the emergence of nitrogenase enzymes as a means to reduce N_2_ to NH_3_ as early as 3.2 billion years ago (Ga).^23^ Today, nitrogenase occurs in a wide array of microbes, from which only three isozymes are known: Mo-, V-, and Fe-nitrogenases, each distinguished by the composition of their active-site metal cofactor.^24–26^ For all three isozymes, two component proteins (dinitrogenase and dinitrogenase reductase) work together to reduce N_2_ to NH_3_, utilizing electrons and MgATP derived from metabolism.^25,26^ Mo-nitrogenase is the most commonly occurring and best studied ^26–29^ and the earliest to evolve.^30^

The essentiality of MgATP in supporting N_2_ reduction by extant Mo-nitrogenase is well known. Intriguingly, divalent metal ions other than Mg^2+^ can support the ATP function (albeit with lower resulting N_2_ reduction rates^31^), yet ATP itself is absolutely required in the mechanism of N_2_ reduction by Mo-nitrogenase, with other nucleotide triphosphates (GTP, UTP, CTP) not supporting N_2_ reduction.^32^ However, considering that generalist function is thought to be a feature of ancient enzymes, it is not known whether the indispensability of ATP is an ancient feature of N_2_ fixation despite the geochemical availability of other cofactors. Specifically, it is unknown whether other nucleoside triphosphates, perhaps coupled to metal ions other than Mg^2+^, may have been capable of supporting an ancient Mo-nitrogenase early in its evolutionary history.

To address this longstanding question, we utilized an experimental paleogenetic approach to investigate the requirement of ATP for ancient biological nitrogen fixation. We reconstructed a Precambrian ancestor of the nitrogenase component proteins, NifH_2_ and NifD_2_K_2_, followed by biochemical and biophysical characterization of the purified ancient proteins with a specific focus on ATP utilization. Our results provide direct laboratory evidence for the necessity of early enzyme cofactor usage of a key enzyme and underscore how biomolecular constraints can predominantly shape metal selection despite far greater availability of metals with similar chemical properties in the environment.

## Results and Discussion

N_2_ reduction by extant Mo-nitrogenase involves the sequential delivery of electrons from the dinitrogenase reductase (NifH_2_) component to the dinitrogenase (NifD_2_K_2_) component, the latter containing the active-site metal cofactor where N_2_ is reduced (**Figure 1**).^26,33^ Each electron delivery event from NifH_2_ to NifD_2_K_2_ requires the hydrolysis of a minimum of 2 MgATP molecules to 2 MgADP and 2P_i_. Under optimal conditions, the minimal reaction stoichiometry can be written as shown in **equation 1**.^29,33,34^

**Figure 1.**
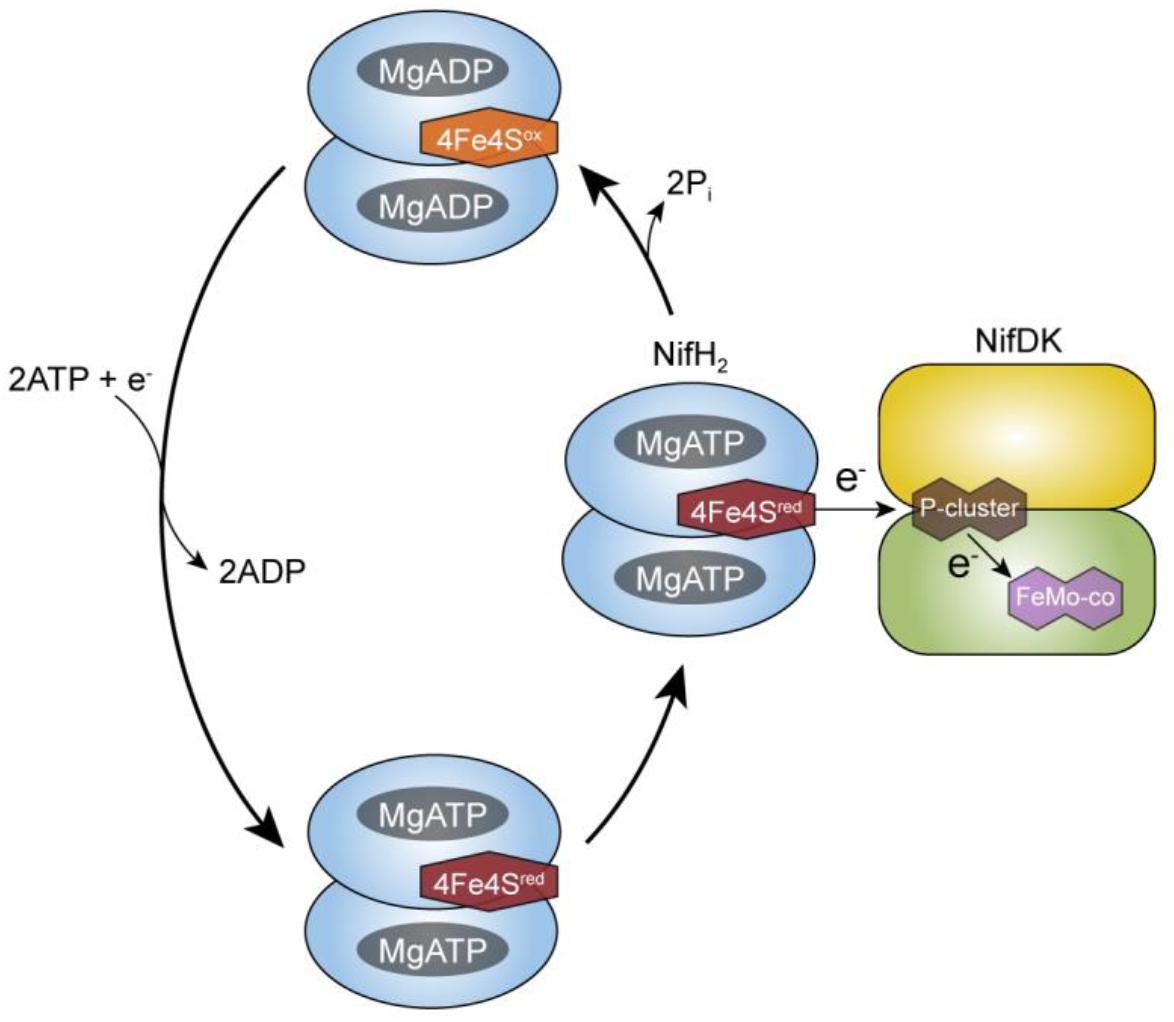
NifH_2_ cycle of Mo-nitrogenase. The NifH_2_ homodimer is shown in blue; it houses two nucleotide binding sites and a [4Fe-4S] redox active cluster. One catalytic half (NifDK) of the NifD_2_K_2_ heterotetramer is shown in yellow (K) and green (D); each catalytic half houses one [8Fe-7S] (P-cluster) and a [7Fe-9S-Mo-C-homocitrate] (FeMo-co) active site metallocluster. When NifH_2_ binds to NifDK an electron is transferred from the [4Fe-4S] cluster to FeMo-co mediated by P-cluster. NifH_2_ then hydrolyzes 2 ATP, dissociates with MgADP bound and an oxidized [4Fe-4S] cluster. MgADP is then exchanged for MgATP and the [4Fe-4S] cluster is reduced, preparing NifH_2_ for binding and electron transfer.

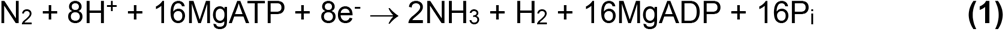

Within this reaction, ATP serves several functions. ATP binds to NifH_2_, inducing conformational changes in the protein that impact its [4Fe-4S] cluster and affinity for binding to NifD_2_K_2_.^27,33,35^ Further, there is growing evidence that the reductase protein induces conformational changes within NifD_2_K_2_ that contribute to substrate binding and reduction.^33,36,37^ Finally, the hydrolysis of ATP and release of the two P_i_ molecules signals the release of NifH_2_ with two bound MgADP from NifD_2_K_2_, with the energy from ATP hydrolysis being used to dissociate the two component proteins.^34,38–40^ The released NifH_2_ is reduced by cellular electron carrier proteins (e.g., ferredoxin or flavodoxin) or non-cellular reductants, such as dithionite.^27^ The two bound MgADP molecules are exchanged with two MgATP molecules, readying NifH_2_ for another round of electron transfer to NifD_2_K_2_ (**Figure 1**).^27,38,39^

We began our study by identifying an ancestral nitrogenase protein (Anc^AK029^) with which to experimentally test ATP utilization in early biological nitrogen fixation (**Figure 2A**). We drew from a prior dataset of ancestral nitrogenase proteins, inferred from a maximum likelihood phylogenetic tree of concatenated NifHDK protein sequences representative of known nitrogenase diversity.^41^ Specifically, we selected the Anc^AK029^ ancestor based on two criteria: having an age well-constrained to the Precambrian and belonging to the direct evolutionary lineage of our laboratory model bacterium *Azotobacter vinelandii* (*A. vinelandii*) (**Figure 2A**). The Anc^AK029^ node also falls within the “Group I” nitrogenase clade primarily hosted by aerobic and facultatively anaerobic bacteria.^41^ Therefore, Anc^AK029^ likely existed after the early oxygenation of the Earth surface environment (Great Oxidation Event, “GOE”) and subsequent diversification of oxygen-tolerant microbes, which provides a maximum age of ∼2.4 Ga.^42^

**Figure 2.**
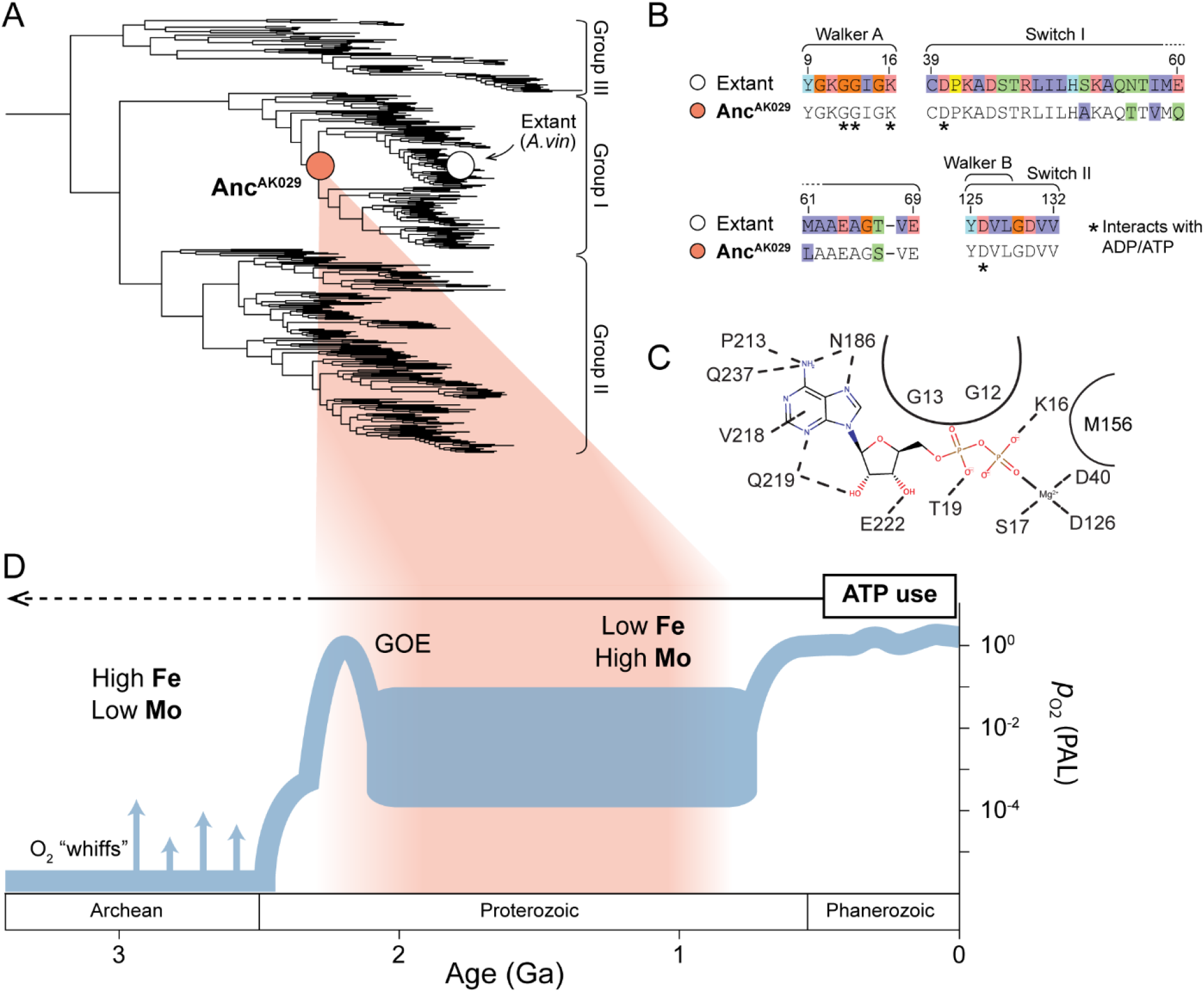
Evolution of the ATP/ADP-binding site in nitrogenase NifH. **(A)** Nitrogenase NifHDK protein phylogeny. Phylogenetic positions of ancestral (Anc^AK029^) and extant *A. vinelandii* (“Extant *A. vin”*) nitrogenase variants discussed in the main text are highlighted by colored circles. **(B)** Protein sequence alignment of NifH ATPase signature motifs (labeled)^33^ Site index from *A. vinelandii* NifH. **(C)** Residue-level intermolecular interactions in the ADP binding site of *A. vinelandii* NifH (PDB 1FP6). All displayed residues are conserved between Anc^AK029^ and WT. Curved lines delineate residues with nonspecific interactions that shape the binding site. **(D)** Estimated age range of nitrogenase ancestors mapped to Earth environmental history. Atmospheric oxygenation plot and relative marine metal abundances are from Lyons et al. (2014).^42^

To gain initial insight into the nucleotide specificity of Anc^AK029^ nitrogenase, we analyzed its global sequence similarity to the Nif nitrogenase of *A. vinelandii* (hereafter simply referred to as the extant nitrogenase), as well as its ancestral features within the inferred nucleotide-binding pocket. Anc^AK029^ NifH, NifD, and NifK proteins together have ∼72% amino acid sequence identity to those of extant nitrogenase. Sequence-level identity is highest between ancestral and extant NifH proteins (∼87%) and lowest between NifK proteins (∼61%). Known ATPase sequence motifs are well conserved in NifH, including the nucleotide-binding P-loop, Walker B motif, and “Switch I and II” motifs (**Figure 2B**).^8,43^ The immediate nucleotide-binding site within NifH includes 14 residues, all of which are identical between Anc^AK029^ and extant nitrogenase (**Figure 2C**). Ten out of these 14 residues are also universally conserved across all extant nitrogenases in our dataset.

### Bacterial growth and biochemical characterization

Expression of ancient, reconstructed enzymes in extant organisms has inherent challenges.^44^ A particular complexity of nitrogenase is the dependence on interactions with multiple protein partners in order to mature a functional enzyme. Thus, a first assessment of the functionality of the inferred ancient proteins is that they can be matured *and* support diazotrophic growth in an extant organism.

Synthetic *nifHDK g*enes encoding the Anc^AK029^ nitrogenase were genomically integrated into the modern nitrogen-fixing model bacterium, *Azotobacter vinelandii*, replacing its native Mo-nitrogenase genes. The results of growth experiments for Anc^AK029^ relative to the extant strain under nitrogen-fixing conditions are shown in **Figure 3**. Growth of the Anc^AK029^ strain is slightly slower overall, but within the error of the extant control, indicating that the ancient nitrogenase is being expressed and is sufficiently functional for N_2_ reduction to support cell growth. Expression was further verified by Western Blot (**Figure S1**), which showed a strong overexpression of the nitrogenase proteins in Anc^AK029^ relative to the extant strain.

**Figure 3.**
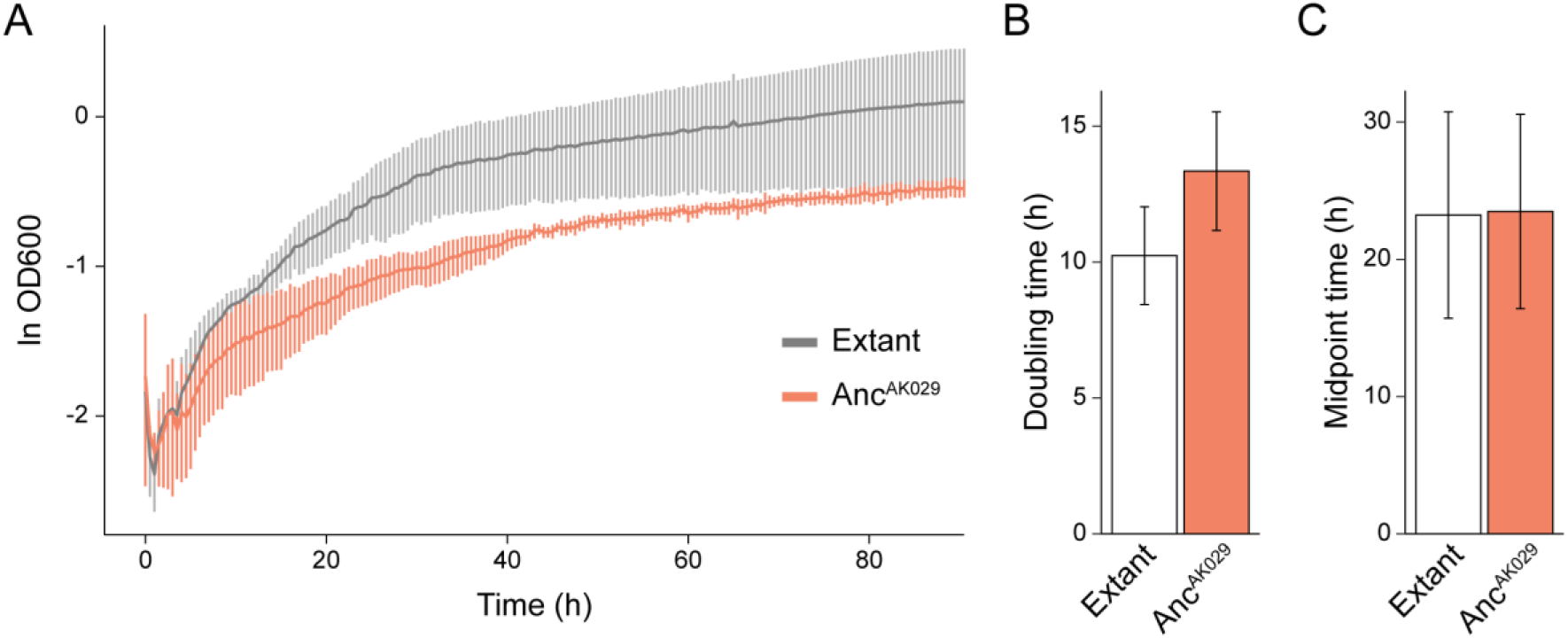
Diazotrophic growth of WT and Anc^AK029^ *A. vinelandii* strains. **(A)** Diazotrophic growth curve of A. vinelandii expressing either the WT or Anc^AK029^ nitrogenase proteins. Line plots represent the average ln OD600 of three biological replicates per strain and error bars indicate ±1 SD. **(B)** Doubling times of WT and Anc^AK029^. **(C)** Midpoint times of WT and Anc^AK029^. **(B, C)** Bars represent the average of three biological replicates per strain and error bars indicate ±1 SD.

Protein purification of Strep-II-NifD_2_K_2_ by streptactin affinity chromatography^45^ and NifH_2_ by anion exchange and size exclusion chromatography^46^ yielded fractions of high purity as determined by SDS-PAGE (**Figure S2**). Densitometry analysis of the SDS-PAGE bands revealed a D:K subunit stoichiometry of 1.44 for Anc^AK029^ versus 0.93 for extant. Fully occupied NifH_2_ contains a [4Fe-4S] cluster with 4 nmol of Fe/nmol NifH_2_. NifD_2_K_2_ contains two FeMo-cofactors (7Fe-9S-Mo-C-homocitrate) and two P-clusters (8Fe-7S) and thus has 2 nmol Mo and 30 nmol Fe/nmol NifDK (Fe:Mo = 15).^24^ Metal analysis by ICP-MS showed Anc^AK029^ NifH_2_ at 2.4 nmol Fe/nmol NifH_2_ (∼60% occupied) and Anc^AK029^ NifD_2_K_2_ at 0.8 nmol Mo/nmol NifD_2_K_2_ and 12.1 nmol Fe/nmol NifD_2_K_2_ (∼40% occupied and Fe:Mo = 14.7). EPR analysis of the [4Fe-4S] cluster of NifH_2_ and FeMo-co of NifD_2_K_2_ in Anc^AK029^ revealed spectra like those established in extant proteins, with a slight perturbation of the FeMo-co g-value (**Figure S3**). Substrate reduction assays showed activities for Anc^AK029^ that are ∼40% of extant MoFe protein for N_2_ reduction under 1 atm N_2_. This decreased activity correlates closely with the metal occupancy and in combination with the densitometry analysis reveals some combination of incomplete maturation or metal loading in the cell. The pure protein is stable over both freeze thaw cycles and when thawed on the bench over the course of multiple assays.

### Alternative nucleotides and divalent metals

The nucleoside triphosphates (e.g., ATP, GTP, ITP, UTP, CTP) used by life are each structurally unique and available in the Precambrian environment,^1^ and they all yield a similar amount of energy upon hydrolysis of the gamma phosphate. Previous studies have shown that extant NifH_2_ can hydrolyze different nucleotides when bound to NifD_2_K_2_, but only ATP supports electron transfer and substrate reduction at NifD_2_K_2_.^32^ These results support the hypothesis that ATP is not simply providing energy for nitrogenase, but that there is something unique to its structure that promotes electron transfer and substrate reduction.^33^ Similar results were found in N_2_ reduction assays here with extant and Anc^AK029^ nitrogenases. No substrate reduction activity or nucleotide hydrolysis was observed for the ancient enzyme with MgGTP, MgITP, and MgUTP (**Figure 4A**). These findings provide evidence for a specific requirement for ATP early in the evolution of nitrogenase.

**Figure 4.**
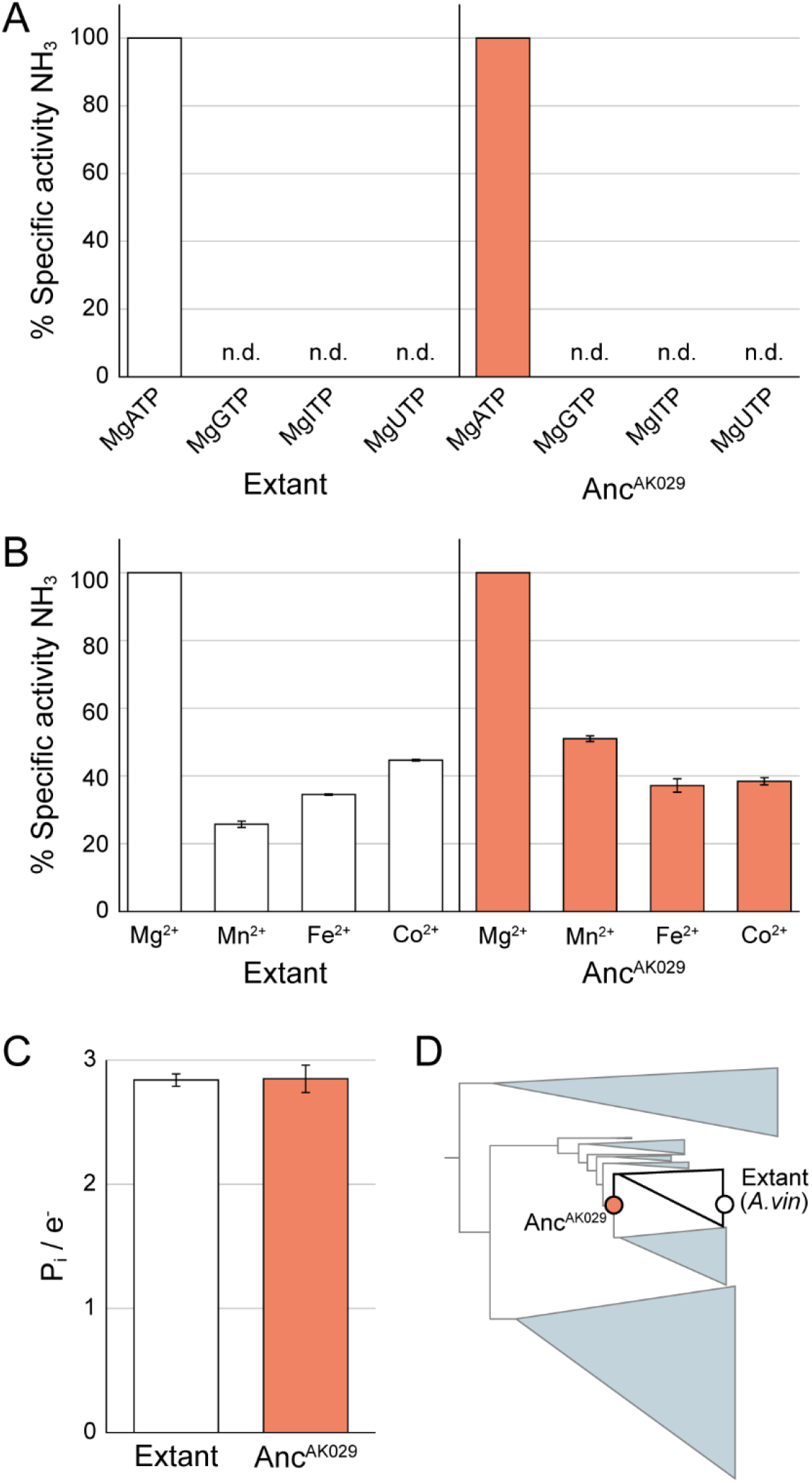
Ancestral nitrogenase behavior with alternative nucleotides and divalent metals. **(A)** Specific activity for N_2_ reduction by extant and Anc^AK029^ NifHDK with different nucleotides complexed with Mg^2+^. Shown are percent specific activities for NH_3_ formation relative to the maximum activity seen with MgATP. n.d., not detectable. **(B)** Specific activity for N_2_ reduction by extant and Anc^AK029^ NifHDK with ATP and different divalent metal ions. Shown are percent specific activities for NH_3_ formation relative to the maximum activity seen with Mg^2+^. Bars represent the average of three replicates and error bars represent ±1 standard deviation. **(C)** Inorganic phosphate released per electron transferred during N_2_ reduction. Shown is inorganic phosphate measured normalized to total electrons in products H_2_ and NH_3_ for extant and Anc^AK029^ nitrogenases when allowed to turn over for 2 min under 1 atm of N_2_. Bars represent the average of three replicates and error bars represent ±1 standard deviation. **(D)** Simplified nitrogenase phylogeny shown in **Figure 2A** indicating evolutionary relationship between variants analyzed in **(A-C)**.

Hydrolysis of ATP by enzymes usually requires a divalent metal ion, facilitating interactions with the enzyme and stabilizing the hydrolysis transition state. The availability of various divalent metal ions has changed throughout Earth’s history, particularly with the oxygenation of the atmosphere.^47^ While many divalent metal ions can serve in the hydrolysis of ATP, Mg^2+^ is the most widely used in extant biological processes and typically supports the highest efficiency and rates.^31,48^ Mg^2+^ is also widely used in many proposed Archaean-aged metabolisms^49^ and has likely remained abundant in the environment,^50^ suggesting that Mg^2+^ would have been available for use by early nitrogen fixing bacteria.

N_2_ reduction assays were performed with different divalent metal ions and ATP for both the extant and Anc^AK029^ nitrogenases (**Figure 4B**). While Mg^2+^ supported the highest activity in both extant and Anc^AK029^, it is not a strict requirement for either enzyme. For example, Mn^2+^, Fe^2+^, and Co^2+^ each showed between 30-50% of the rates of substrate reduction in the Anc^AK029^ nitrogenase compared to the rates with Mg^2+^ (**Figure 4B**). These findings reveal that while Mg^2+^ provides the highest specific activities for N_2_ reduction for both extant and the Anc^AK029^ enzymes, other divalent metals can also support activity and could be utilized depending on the metals available in the environment.

### ATP/e^-^ usage

Extant NifH_2_ hydrolyzes ∼2 ATP *only* while complexed with NifDK, which is coupled to the transfer of a single electron per association event (**Figure 1**).^33,34,38^ This ratio of ATP/e^-^ is a marker of the specificity and control of the system. While NifH_2_ has all the makings of a canonical ATPase,^33,51^ only the full NifH_2_-NifDK complex shows any detectable ATPase activity. This phenomenon allows for efficient use of costly ATP and reducing equivalents and has been suggested to involve mechanisms that promote N_2_ binding and reduction.^36,37^ To test the linkage of MgATP hydrolysis to electrons transferred to substrates, the inorganic phosphate released during N_2_ reduction assays was measured to quantify the ATP hydrolyzed. The amount of ATP hydrolyzed was compared to total electrons transferred by totaling all electrons in the products H_2_ and NH_3_. As can been seen in **Figure 4C**, Anc^AK029^ nitrogenase shows a P_i_/e^-^ ratio identical to that measured for extant nitrogenase of ∼2.8 ATP/e-, which is in agreement with prior studies on extant nitrogenase.^34^ These results are further evidence for the tight linkage between MgATP hydrolysis and substrate reduction for both Anc^AK029^ and extant nitrogenases.

## Summary

Energy transfer to drive enzymatic reactions is a fundamental feature of life. However, understanding of the evolution of the cellular suite of energy carrying molecules has been stalled by the lack of investigative strategies to directly test the nucleotide triphosphate specificities of key ancient metabolic enzymes. Paleogenetic approaches like that employed in the present work enable functional insights that are not possible by examining extant enzymes alone. Ancestral sequence reconstruction has previously been used to investigate the evolutionary drivers of structural complexity,^52^ the origins of novel enzymatic activity and specificity,^53^ and ancient biosignature generation by geobiologically critical enzymes.^54^ Recently, ancestral nitrogenase enzymes were resurrected, providing evidence of a long-term conservation of the N_2_-binding mechanism in biological nitrogen fixation.^41^ Now, our finding that the strict requirement for ATP by nitrogenase extends back to the Precambrian not only addresses this gap by providing direct evidence of ancient biological ATP usage, but underscores its essential role for a microbial metabolism that has limited the biosphere billions of years.^47,55^

Extant Mo-nitrogenase demonstrates a unique relationship with ATP. The system not only requires the energy from hydrolysis of ATP, but the binding of ATP itself is a strict mechanistic requirement for electron transfer from NifH_2_ to NifD_2_K_2_ and the ability to bind and reduce N_2_. Further, while NifH_2_ has all the canonical sequence motifs of an ATPase, it is only the NifH_2_-NifD_2_K_2_ complex that hydrolyzes 2 ATP coupled to the transfer of a single electron. In this way, the system exerts tight control over the use of valuable ATP and reducing equivalents. Here, it is demonstrated that these specific aspects of the ATP-dependent mechanism of nitrogenase likely emerged early in the evolution of this enzyme. The reconstructed Anc^AK029^ shows strong conservation of ATP relevant sequence motifs in NifH_2_, supports diazotrophic growth in *A. vinelandii*, and the purified enzyme is stable and active for N_2_ reduction. Like extant nitrogenase, N_2_ reduction activity in Anc^AK029^ is seen exclusively with ATP; alternative nucleotides (GTP, UTP, ITP) are neither hydrolyzed nor support catalysis in the system. ATP hydrolyzed per electron transferred in Anc^AK029^ is identical to extant nitrogenase at ∼2.8 ATP/e. Finally, a number of divalent metals (Mn^2+^, Fe^2+^, Co^2+^) are shown to support the function of ATP, but none approach the rates supported by Mg^2+^.

What is notable is the specificity of MgATP to the Mo-nitrogenase mechanism and, as evidenced here, how early this relationship formed, particularly given that the activities of other ATPases can be supported by other nucleoside triphosphates.^10^ The complexity, specificity, and control of the Mo-nitrogenase mechanism are testaments to the challenge of N_2_ reduction and the early evolution of these features speaks to the demand for a suitable catalyst by early life, and the unique role that ATP played in meeting that demand.

## Associated Content

Additional data are available in the Supporting information (supplemental figures, tables, methods, and materials). All unique biological materials (i.e., *Azotobacter* strains) will be available upon request.

## Protein Accession IDs

Extant nitrogenase molybdenum-iron protein alpha chain – NifD, UniProtKB P07328 Extant nitrogenase molybdenum-iron protein beta chain – NifK, UniProtKB P07329 Extant nitrogenase iron protein 1 – NifH, UniProtKB P00459

## Author Information

### Corresponding Authors

L.C.S.

B.K.

## Author contributions

AKG and BK performed the ancestral sequence reconstruction. SDC generated the Anc^AK029^ *A. vinelandii* strain. DFH performed N_2_ reduction assays for alternative nucleotides and divalent metal ions. HRR performed growth experiments and Western Blot analysis. ZYY performed the ATP hydrolysis experiments and EPR analysis. HF generated the metals analysis, SDS-PAGE, and densitometry analysis. DFH, LCS, HRR, AKG, and BK wrote the manuscript.

## Notes

The authors declare no competing financial interest.

## Acknowledgements

This work was supported by the National Aeronautics and Space Administration (NASA) Interdisciplinary Consortium for Astrobiology Research: Metal Utilization and Selection Across Eons, MUSE 80NSSC22K0546 (PI: Kaçar). We thank Dennis Dean and Valerie Cash for providing *A. vinelandii* strains DJ and DJ2102; and the members of the Metal Selection and Utilization Across Eons (MUSE) ICAR for helpful discussions.

