## Supplemental Information for "The Origins of ATP Dependence in Biological Nitrogen Fixation"

### Supplemental figures

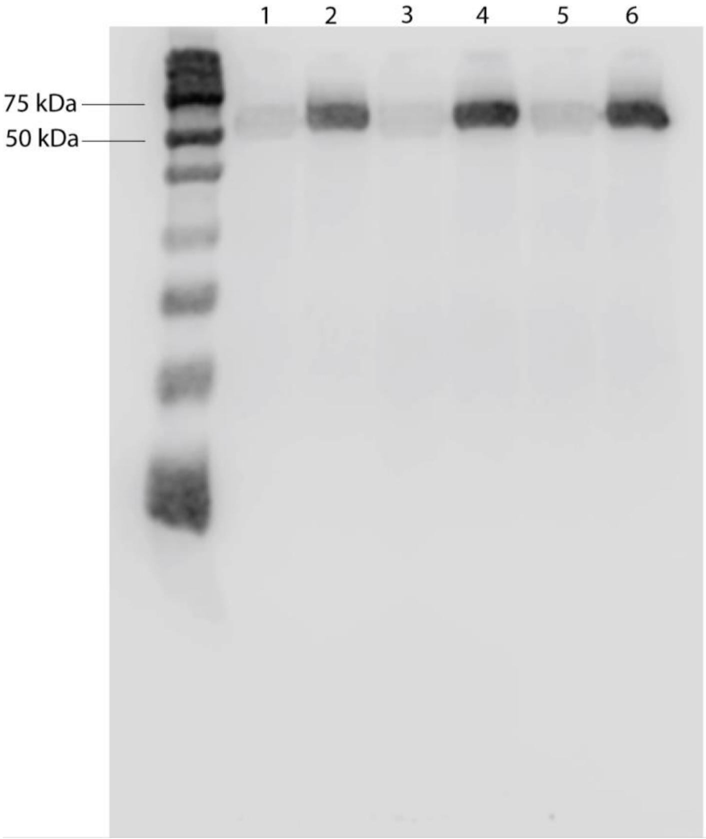


**Figure S1**. Immunodetection of Strep-II-tagged NifD in WT (lanes 1, 3, and 5) and Anc *A. vinelandii* strains (lanes 2, 4, and 6). Detection of Strep-II-tagged NifD was determined using an anti-Strep antibody. Each lane represents one biological replicate (three replicates per strain). The expected size of NifD is 55.8 kDa.

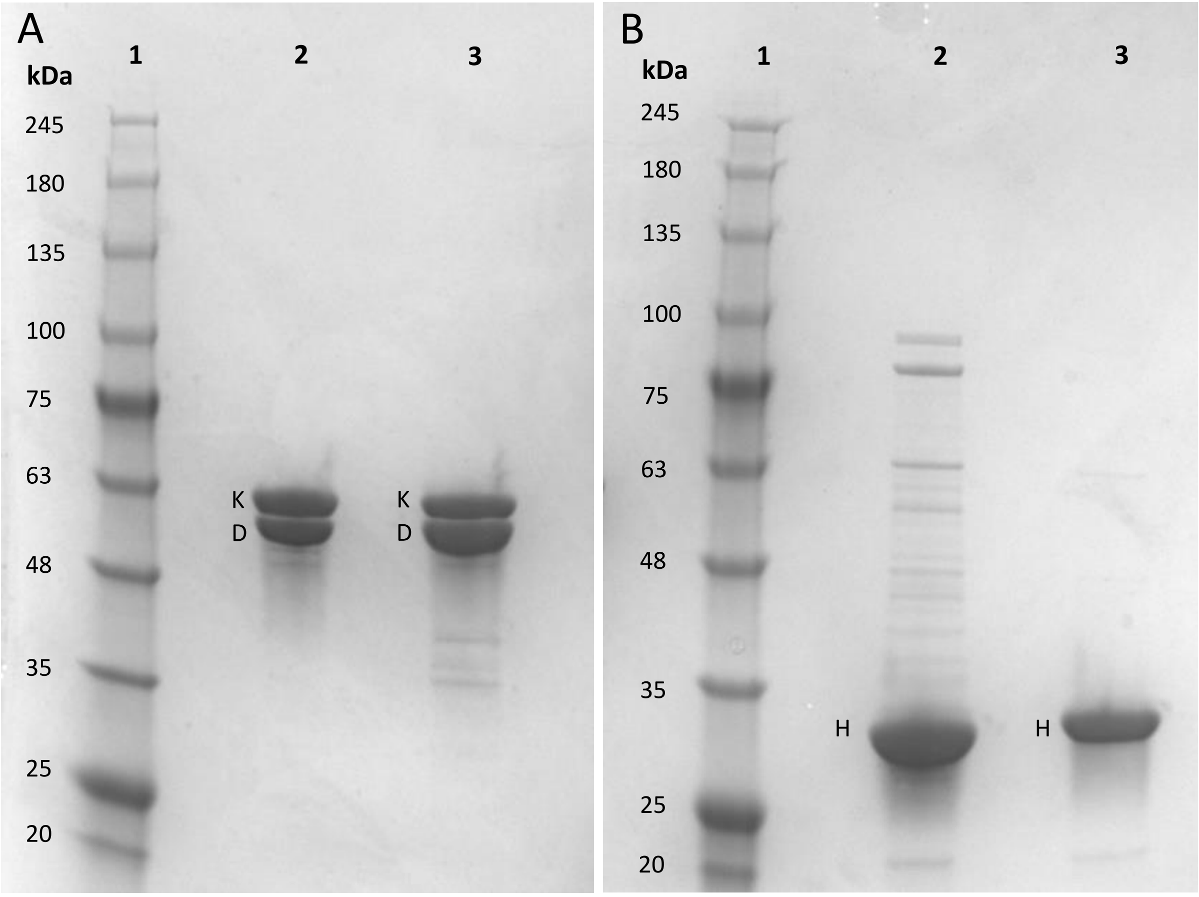


C

| | Intensity | Molecular Weight (kDa) | Intensity/kDa | Stoichiometry (D:K) |
|---|---|---|---|---|
| Extant K | 19,785,513 | 57,530 | 343.92 | 0.93 |
| Extant D | 17,344,927 | 54,230 | 319.84 | |
| Anc K | 18,160,805 | 56,100 | 323.72 | 1.44 |
| Anc D | 24,734,047 | 53,020 | 466.50 | |

**Figure S2.** Coomassie stained SDS-PAGE and densitometry analysis. A) Shown is protein ladder (lane 1) and purified Extant (lane 2) and Anc (lane 3) NifD and NifK subunits. B) Shown is protein ladder (lane 1) and purified Extant (lane 2) and Anc (lane 3) NifH. C) Densitometry analysis of NifDK.

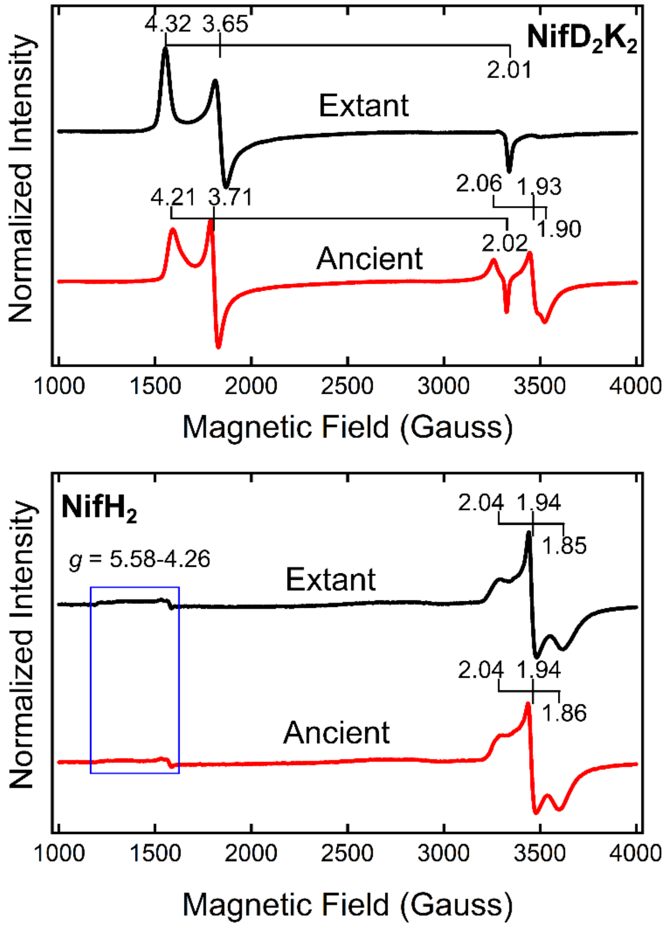


**Figure S3.** Continuous-wave (CW) EPR spectra of extant (black traces) and ancient (red traces) $NifD_2K_2$ (top panel) and $NifH_2$ (bottom panel) proteins in the dithionite-reduced resting state. The EPR spectra were recorded from samples containing 50 μM of each protein at 12 K and 20 mW microwave power. Each spectrum is a sum of 5 scans. The ancient $NifD_2K_2$ displayed a similar $S$ = 3/2 spin signal as that in the extant $NifD_2K_2$ with slight difference in rhombicity as indicated by the $g$ values (top panel). Moreover, an $S$ = 1/2 signal in the $g \sim 2$ region (g = 2.06, 1.93, and 1.90) might originate from an immature P-cluster species.[1] The ancient $NifH_2$ exhibited almost identical EPR features as those of the extant $NifH_2$, with the blue box highlighting the low intensity high spin states (bottom panel).

### Methods and Materials

*Ancestral sequence reconstruction*. Nitrogenase phylogenetic construction and ancestral sequence reconstruction was performed as described by Garcia et al. 2020, 2022, 2023.[2–4] Briefly, a nitrogenase protein sequence dataset was curated by BLASTp[5] against the NCBI non-redundant protein database (accessed August 2020) using *A. vinelandii* NifH, NifD, and NifK query sequences. Sets of homologs for each nitrogenase subunit were aligned by MAFFT v7.450[6] together with outgroup dark-operative protochlorophyllide oxidoreductase sequences (Bch/ChlLNB) and concatenated into a single alignment. Phylogenetic reconstruction by RAxML v8.2.10 was performed using the untrimmed, concatenated alignment and best-fit model parameters, LG+G+F, assessed by ModeFinder[7] implemented in IQ-TREE v.1.6.12[8]. Clade support was evaluated by the SH-like aLRT (Anisimova and Gascuel, 2006). Ancestral sequence reconstruction was performed by PAML v4.9j[9] using the same model parameters used for phylogenetic reconstruction, generating the ancestral nitrogenase NifHDK protein sequence "Anc$^{AK029}$" targeted in the current study (see Figure 2A). Nitrogenase phylogenies were visualized using the ggtree v.3.8.0 R package.

*A. vinelandii strain engineering*. Strains and plasmids used in the present study are listed in **Table** S1. The most probable ancestral NifHDK protein sequence (i.e., the sequence with the most probable ancestral residue per site) reconstructed for node Anc$^{AK029}$ was reverse-translated and codon-optimized for expression in *A. vinelandii* ("wild-type", WT), as described by Garcia et al. 2023.[2] Sites for which the Anc$^{AK029}$ and WT amino acid residues were identical were assigned the WT codon, and sites for which the ancestral and WT amino acid residues differed were assigned a random codon weighted by *A. vinelandii* codon frequencies (Codon Usage Database, https://www.kazusa.or.jp/codon/as). The optimized Anc$^{AK029}$ *nifH*, *nifD*, and *nifK* nucleotide sequences were concatenated together with *A. vinelandii* intergenic regions and 1000-bp flanking sequences to direct homologous recombination at the native *nifHDK* locus in the *A. vinelandii* genome. In addition, an "ASWSHPQFEK" Strep-II tag was appended to the N-terminus of ancestral NifD for downstream affinity purification. The concatenated sequence was synthesized and cloned into pUC19 (GenScript, Piscataway, NJ, USA), yielding plasmid pAnc$^{AK029}$.

Genomic integration of ancestral *nifHDK* into *A. vinelandii* followed Garcia et al. 2023,[2] using established methods for transformation and homologous recombination.[10] Genetically competent cells were prepared from the non-diazotrophic *A. vinelandii* parent strain, "Δnif", containing disruptions to all three *nif*, *vnf*, and *anf* nitrogenase gene clusters,[11] by growth in Mo- and Fe-deficient BN medium. The Anc$^{AK029}$ strain harbors a clean replacement of *nifHDK* with a kanamycin resistance cassette (KanR). Competent cells were transformed by congression (i.e., the coincident introduction of two unrelated genetic modifications)[10] using ~1 µg of plasmid pDB303 and 1 µg of digested (ScaI-HF, New England BioLabs) plasmid pAnc$^{AK029}$, the former containing the rifampicin resistance determinant *rpoB113* (RifR). Transformants were screened for rifampicin resistance and loss of kanamycin resistance and confirmed via both Sanger sequencing (primers listed in **Table S1**) and Oxford Nanopore whole-genome sequencing (Plasmidsaurus, Eugene, OR, USA). Transformants were stored at -80℃ in phosphate buffer with 7% DMSO.

*A. vinelandii growth*. Cultures were grown in liquid or solid Burk's media ("BN medium") with 1 μM $Na_2MoO_4$ and 10 mM ammonium acetate. To induce diazotrophic growth, Burk's media without ammonium acetate was also used ("B medium"). 50 mL seed cultures in BN medium were grown for 24 hours at 30°C with a 300 rpm double orbital agitation for nitrogenase expression and growth rate quantification.

Growth rate analysis followed the protocol described in Carruthers et al.,[12] as summarized below. Three biological replicates of Anc$^{AK029}$ and WT strains were inoculated from the 24-hour seed cultures into liquid B medium at an optical density at 600 nm (OD600) of 0.05. The cultures were aliquoted into a 96-well, flat-bottom plate (Greiner Bio-One) and sealed with a Breathe-Easy adhesive membrane (Diversified Biotech). The plate was placed in a SPECTROstar Nano Microplate Reader ((BMG Labtech, Ortenberg, Germany) and incubated at 30°C with a 300 rpm double orbital agitation. The OD600 of the cultures was measured every 30 minutes for 90 hours. The R package Growthcurver was used to calculate doubling time and visualize data.[13] Statistical significance was determined using one-way ANOVA and post-hoc Tukey HSD test.

*Protein quantification*. Protein quantification of Strep-II-tagged NifD was conducted for Anc$^{AK029}$ and DJ2102 (containing a Strep-II-tagged WT NifD) strains following the protocol in Garcia et al., 2023[2] with modifications as described below. Flasks containing 100 mL B medium were inoculated to an OD600 of 0.01 and grown diazotrophically. Cell pellets were harvested after 4 hours and stored at -80°C. TE lysis buffer (10 mM Tris, 1 mM EDTA, 1 mg/mL lysozyme) was prepared and used to resuspend the cell pellets. The resuspended cell pellets were vortexed lightly and heated for 10 minutes at 95°C.The lysates were centrifuged for 15 minutes at 5000 rpm. The supernatants were analyzed for total protein quantification using the Pierce BCA Protein Assay kit (ThermoFisher). The lysates were normalized to 200 μg total protein and diluted 1:1 with 2x Laemmli buffer before performing polyacrylamide gel electrophoresis (PAGE). The proteins from the PAGE gel were transferred to a nitrocellulose membrane (ThermoFisher). The membrane was stained with Revert 700 Total Protein Stain (LI-COR) and imaged with an Odyssey Fc Imager (LI-COR). The membrane was destained using Revert Destaining Solution (LI-COR) and blocked for 1 hour using 5% non-fat milk dissolved in PBS (137 mM NaCl, 2.7 mM KCl, 10 mM $Na_2HPO_4$, 1.8 mM $KH_2PO_4$). The membrane was rinsed with a PBS and 0.01% Tween-20 mixture (PBS-T) to remove residual blocking solution. The membrane was then incubated with primary Strep-II antibody (Strep-MAB-Classic, IBA Lifesciences, 1:5000 in 0.2% BSA) and gently rocked at 10°C for 16 hours and then at room temperature for 2 additional hours. The membrane was rinsed with PBS-T and incubated with 1:15,000 IRDye 680RD Goat anti-Mouse in LI-COR blocking buffer for 2 hours at room temperature. The membrane was then imaged again using an Odyssey Fc Imager (LI-COR).

*Purification reagents and procedures.* Reagents were obtained from Sigma-Aldrich (St. Louis, MO) or Fisher Scientific (Fair Lawn, NJ) and used without further purifications. Argon and dinitrogen gases were purchased from Air Liquide America Specialty Gases (Plumsteadville, PA). All manipulations of proteins and buffers were done anaerobically in septum-sealed serum vials and flasks utilizing a vacuum Schlenk line under argon or dinitrogen atmospheres and gas-tight syringes.

*Protein purification and quantification.* $NifD_2K_2$ and $NifH_2$ proteins were expressed and purified from *Azotobacter vinelandii* strains DJ2102 (Extant Strep-$NifD_2K_2$), DJ884 (Extant $NifH_2$), $Anc^{AK029}$ (Ancient Strep-$NifD_2K_2$ and $NifH_2$) by previously published methods.[14,15] Protein concentration was quantified by the Biuret method[16] with bovine serum albumin as a standard and purity assessed at ≥ 90% by SDS-PAGE with Coomassie blue staining. The $H^+$ reduction activity of extant $NifD_2K_2$ was assessed at 8.5 nmol $H_2$/nmol $NifD_2K_2$/s, which is within the established range for fully active $NifD_2K_2$ from *Azotobacter vinelandii*.[17–20] The $H^+$ reduction activity of $Anc^{AK029}$ $NifD_2K_2$ was assessed at 3.8 nmol $H_2$/nmol $NifD_2K_2$/s.

*EPR.* EPR samples of extant and Anc $NifD_2K_2$ and $NifH_2$ were prepared with 50 µM of each protein. $NifH_2$ was prepared in a buffer of 50 mM Tris pH 7.5, 100 mM NaCl, and 10 mM dithionite. $NifD_2K_2$ was prepared in a buffer of 100 mM MOPS pH 7, 5 mM ATP, 10 mM phosphocreatine, 7.5 mM $MgCl_2$, 0.2 mg/mL creatine phosphokinase, and 1.3 mg/mL BSA. Prepared samples were frozen in a pentane/liquid $N_2$ slurry. Continuous-wave (CW) X-band EPR spectra were recorded using a Bruker ESP-300 spectrometer with an EMX PremiumX microwave bridge and an $EMX^{PLUS}$ standard resonator in perpendicular mode, equipped with an Oxford Instruments ESR900 continuous helium flow cryostat using VC40 flow controller for helium gas. Spectra were recorded at the following conditions: temperature, ~12 K; microwave frequency, ~9.38 GHz; microwave power, 20 mW; modulation frequency, 100 kHz; modulation amplitude, 8.14 G; time constant, 20.48 ms. Each spectrum is the sum of five scans.

*Substrate reduction assays.* Assays were performed in 9.4 mL vials with a nucleotide regeneration buffer (6.7 mM $MgCl_2$, 30 mM phosphocreatine, 5 mM ATP, 0.2 mg/mL creatine phosphokinase, 1.2 mg/mL BSA) and the reductant 10 mM sodium dithionite in 100 mM MOPS buffer at pH 7.0. Reaction vials were made anaerobic and put under an atmosphere of $N_2$ gas. In assays that tested different nucleoside triphosphates (GTP, ITP, UTP), the ATP in the buffer was replaced with the appropriate nucleoside triphosphate at the same concentration. Controls with no nucleoside triphosphate added had no activity. In assays that tested alternative divalent metal ions ($Mn^{2+}$, $Fe^{2+}$, $Co^{2+}$), the $Mg^{2+}$ was replaced with the appropriate metal at the same concentration. Controls with no metal added had no activity. All assays were performed with 0.42 µM $NifD_2K_2$ and 8.4 µM $NifH_2$. $NH_3$ was quantified using a fluorescence protocol[21] with some modifications. An aliquot of the sample was added to a solution containing 200 mM potassium phosphate pH 7.3, 20 mM o-phthalaldehyde, and 3.5 mM 2-mercaptoethanol and incubated for 30 minutes in the dark. Fluorescence was measured at $\lambda_{excitation}$ of 410 nm and $\lambda_{emission}$ 472 nm and

$NH_3$ was quantified using a standard generated with $NH_4Cl$. $H_2$ was measured and quantified using a molecular sieve 5A column and thermal conductivity detector.

*ATP hydrolysis.* Assays were performed in a reaction buffer with 10 mM MgATP and 10 mM dithionite in 100 mM MOPS buffer at pH 7.3, without a MgATP regeneration system. The protein concentrations were 0.42 µM $NifD_2K_2$ and 8.4 µM $NifH_2$. The ratios of hydrolyzed ATP per electron transferred for product formation under $N_2$ were determined by quantification of total amount of $P_i$ versus total amount of electrons in the products $H_2$ and $NH_3$ as previously described.[22]

**Table S1. Strains, plasmids, and primers used in the current study.**

| Type | Designation | Source | Additional Information |
|---|---|---|---|
| *A. vinelandii* strain | WT (DJ) | DOI:10.1128/JB.00504–09 | Dennis Dean, Virginia Tech; Wild-type (WT); Nif+ |
| *A. vinelandii* strain | DJ2102 | DOI:10.1016/bs.mie.2018.10.007 | Dennis Dean, Virginia Tech; Strep-tagged WT NifD; Nif+ |
| *A. vinelandii* strain | Δnif | Russell et al., 2024 | Δ*nifHDK*::KanR + Δ*vnfDGK*::StrR + *anfD*::GenR; Nif-, Vnf-, Anf- |
| *A. vinelandii* strain | Anc$^{AK029}$ | This paper | Δ*nifHDK*::*nifHDK*$^{Anc}$ (Strep-tagged NifD) + Δ*vnfDGK*::StrR + *anfD*::GenR; Nif+, Vnf-, Anf- |
| plasmid | pDB303 | Dennis Dean | 1.7-kbp EcoRI *A. vinelandii* fragment containing RifR determinant rpoB113 |
| plasmid | pAnc$^{AK029}$ | This paper | *nifHDK*$^{Anc}$ (Strep-tagged NifD) + 1000-bp *nifHDK* homology sequences , in pUC19 vector |
| primer | 306_nifH_F | DOI:10.7554/eLife.85003 | GCCGAACGTTCAAGTGGAAA |
| primer | 307_nifH_R | DOI:10.7554/eLife.85003 | AGAGCCAATCTGCCCTGTC |
| primer | 308_nifD_F | DOI:10.7554/eLife.85003 | CACCCGTTACCCGCATATGA |
| primer | 309_nifD_R | DOI:10.7554/eLife.85003 | ACTCATCTGTGAACGGCGTT |
| primer | 310_nifK_F | DOI:10.7554/eLife.85003 | GCTAACGCCGTTCACAGATG |
| primer | 311_nifK_R | DOI:10.7554/eLife.85003 | TCAGTTGGCCTTCGTCGTTG |
